# Macrophage Antigen Presentation Is Unleashed by Pan-RAS Inhibition to Promote Antitumor Immunity

**DOI:** 10.64898/2026.01.30.702886

**Authors:** Jie Shen, Weini Li, Xin He, Lianjun Zhang, Huidi ShuCheng, ChunBo Zeng, Mingtian Che, Wei Fan, Suda Jo, Heping Yang, Yuan Yuan, Zakir Khan, Zhi Chen, DuoYao Cao

## Abstract

Pan-RAS inhibitors have emerged as potent targeted therapies designed to suppress oncogenic signaling in RAS-mutant tumors. However, whether these inhibitors possess therapeutic utility beyond tumors with intrinsic RAS dependency remains unclear. Here we show that the clinical-stage pan-RAS(ON) inhibitor RMC-6236 (Daraxonrasib) induces robust regression of NRAS–wild-type B16 tumors in immunocompetent mice, a response mediated by a mechanism independent of direct tumor-cell targeting. Instead, RMC-6236 acts by remodeling the tumor microenvironment, specifically enhancing the antigen-presenting capacity of tumor-associated macrophages through upregulated MHC-I expression. This anti-tumor efficacy is abrogated in T cell-deficient hosts, identifying a macrophage-dependent, T cell-mediated mechanism of control. Mechanistically, RMC-6236 does not directly block canonical RAS signaling in macrophages. Rather, it suppresses a Myc-Hdac2 epigenetic program, leading to enhanced histone H4 acetylation at the *Nlrc5* locus and subsequent activation of the MHC-I master regulator NLRC5. Together, our findings reveal a macrophage-centered, epigenetically wired immune mechanism for pan-RAS inhibition that broadens the therapeutic scope of targeting the “undruggable” RAS pathway beyond tumor genotype towards systemic immune modulation.

## Introduction

Tumor-associated macrophages (TAMs) are a dominant immunosuppressive population within the tumor microenvironment (TME) that actively dampens anti-tumor immunity by promoting tumor growth and therapeutic resistance. Rather than functioning as effective innate immune sentinels, TAMs exhibit impaired phagocytic activity and dysfunctional antigen presentation, thereby blunting productive T-cell responses and facilitating tumor progression1,2. Accordingly, therapeutic strategies that reprogram TAMs toward an immune-activated, anti-tumor phenotype represent a promising avenue to enhance host anti-tumor immunity.

In addition to cytokine- and metabolite-mediated cues within the TME, intracellular oncogenic signaling pathways traditionally studied in tumor cells are increasingly recognized as critical regulators of immune cell function. Among these, the RAS oncogene family (HRAS, KRAS, and NRAS) constitutes one of the most frequently mutated drivers in human cancers ^3,4^. Beyond their cell-intrinsic oncogenic functions, accumulating evidence indicates that oncogenic KRAS, as a prototypical RAS oncoprotein, profoundly shapes the tumor immune landscape by promoting macrophage recruitment, expansion, and polarization toward an immunosuppressive TAM phenotype ^5-7^. These observations raise the possibility that therapeutic inhibition of RAS signaling may influence not only tumor cells but also immune cells within the TME.

Recently, the RAS(ON) multi-selective inhibitor RMC-6236 (Daraxonrasib) has demonstrated robust tumor-suppressive activity across multiple RAS-mutant cancers in preclinical models ^8^. Intriguingly, emerging studies suggest that RAS inhibition not only alters tumor cell signaling but also remodels the immune microenvironment, including increased macrophage infiltration and enhanced CD8^+^ effector T-cell–mediated immunity ^9-12^. Nevertheless, whether RMC-6236 exerts direct immunomodulatory effects on TAMs themselves remains poorly understood.

Although RMC-6236 is a pan-RAS inhibitor theoretically capable of targeting multiple RAS isoforms, prior studies have predominantly focused on KRAS-driven cancer models, leaving its therapeutic potential in RAS–wild-type tumors poorly defined^13,14^ [38589574, 39586491]. B16 melanoma is wild-type for RAS [14743208]^15^, thereby providing an experimental context in which the effects of pan-RAS inhibition cannot be attributed to direct suppression of oncogenic RAS signaling within tumor cells. In this setting, we observed that RMC-6236 exerted minimal tumor-intrinsic effects in vitro but induced pronounced tumor suppression in vivo in immune-competent hosts, strongly implicating tumor-extrinsic, immune-mediated mechanisms.

Accordingly, this model enables focused interrogation of how RMC-6236 modulates macrophage function within the TME, permitting mechanistic dissection of macrophage-centered immune regulation underlying pan-RAS inhibition.

## Materials and Methods

### Drugs and reagents

A complete list of inhibitors and reagents used in this study, including vendor information and catalog numbers, is provided in **Supplementary Table 1**.

### Animal experiments

All male and female mice were fed PicoLab® Rodent Diet 20 (5053; Newco Specialty), the standard chow used by our animal facility. WT and Nude (**NU/J**) Mice were inoculated subcutaneously (s.c) in the skin on their backs with 5×10 ^5^ B16-F10-OVA or B16-F10 melanoma cells suspended in 100 µL PBS. Tumor volume was measured every 4 days, the formula: V = (Length (L) × Width (W)^2^) × 0.52, where L > W. Starting on day 7, the mice were daily gavage RMC-6236 treatment (10mg/KG) as previous suggested ^8^. Also, we collected the facial blood every 4 days to analyze the monocytes. On day 21, the mice were sacrificed, and macrophages and CD8^+^ T cells were analyzed.

#### Ficoll Gradient Isolation of Mouse PBMCs

Peripheral blood mononuclear cells (PBMCs) were isolated from mouse blood using Lymphoprep™ (STEMCELL Technologies), a density gradient medium functionally equivalent to Ficoll-Paque. Briefly, 300–400 μL of blood was collected into EDTA-coated tubes and diluted 1:1 with 1× PBS containing 2% FBS (Omega, Cat# FB-12). The diluted blood was gently layered over 2 mL of Lymphoprep™ in a 5 mL polystyrene tube, taking care not to disturb the interface. Samples were centrifuged at 800 × g for 20 minutes at room temperature with the brake off. The PBMC-enriched layer at the plasma–Lymphoprep interface was carefully aspirated using a 200 μL pipette tip. Isolated PBMCs were immediately processed for surface staining and analysis by flow cytometry ^16^.

#### Thioglycolate-Induced peritoneal Macrophages (TPM)

Mice were intraperitoneally injected with thioglycolate four days prior to sacrifice to induce the recruitment of peritoneal macrophages. Briefly, melanoma-bearing mice were intraperitoneally injected with sterile thioglycolate broth four days prior to sacrifice (day 17 post-tumor implantation). Mice were sacrificed on day 21, macrophages were isolated by peritoneal lavage using medium B, which consisted of HBSS supplemented with penicillin (100 U/mL), streptomycin (100 μg/mL), 10 mM HEPES (pH 7.4), 0.06% BSA, and 0.0375% NaHCO_3_, as previously described ^17^. The isolated cells were cultured in medium C: RPMI 1640 (without L-glutamine) supplemented with 10% FBS, 1 U/mL penicillin, 1 μg/mL streptomycin, 0.5 mM sodium pyruvate, 10 mM HEPES, 50 μM 2-mercaptoethanol, and 1 ng/mL M-CSF (PeproTech). After a 4-hour adherence period, non-adherent cells were removed by washing the plates three times with PBS as previously described ^17^

#### Tumor-Associated Macrophage Isolation

Melanoma tissues were harvested and processed into single-cell suspensions. Macrophages were then isolated using F4/80 magnetic microbeads (Miltenyi Biotec, Cat# 130-110-443) according to the manufacturer’s instructions. The cell suspensions were passed through MS Columns (Miltenyi Biotec, Cat# 130-042-201) to enrich for F4/80^+^ cells. The isolated macrophages were subsequently used for downstream analyses and experiments.

#### Monocyte/macrophage depletion

To deplete monocytes/macrophages, mice were treated with 300 μg anti-CSF1R antibody (clone AFS98, Bio X Cell, Cat. No. BP0213) via intraperitoneal injection every 3 days for the indicated pre-treatment period before tumor challenge ^18,19^. Control animals were injected with an equivalent amount of rat IgG2a isotype control (Bio X Cell, Cat. No. BP0089).

All animals were housed under a 14-hour light / 10-hour dark cycle, with lights on from 6:00 AM to 8:00 PM. The Rodent room temperatures are controlled at 74 ± 2°F. All animal procedures were approved by IACUC at Cedars-Sinai Medical Center and conducted in accordance with the NIH Guide for the Care and Use of Laboratory Animals. Additional experiments performed in China were approved by the IACUC at Hainan Medical University. Mice were euthanized by cervical dislocation following anesthesia with an overdose of isoflurane. All efforts were made to minimize animal suffering and ensure ethical treatment throughout the study.

### Cell Culture and treatment

#### Macrophages phagocytosis

B16 melanoma cells (0.5 × 10^6^) cells were treated with 5 µM CFSE (Biolegend) for 30 minutes, followed by three washes with PBS. The CFSE-labeled melanoma cells were then fed to melanoma-associated macrophages for 12 hours. After this incubation, the phagocytic ability of the macrophages was assessed by measuring the CFSE signal within the macrophages. The percentage of macrophages that had taken up melanoma cells was assessed using a Sony ID7000 flow cytometer (Sony). The data were analyzed using FlowJo software version 10.4 (FlowJo, LLC, Ashland, OR).

#### Macrophages antigen uptake and presentation

Melanoma-associated macrophages (1.0 × 10^6^/mL) were incubated with Alex 647-labeled OVA (OVA-Alex 647; 500 ng/mL) (Thermo) in complete RPMI medium at 37°C for 2 hours. After incubation, the cells were washed with PBS containing 2% FBS and analyzed by flow cytometry. The mean fluorescence intensity (MFI) from internalized OVA-Alex 647 was used to evaluate the macrophages’ antigen uptake ability. Additionally, H-2Kb-restricted OVA (SIINFEKL) MHC class I peptides were fed to the tumor-associated macrophages for 24 hours. Following this incubation, the cells were washed three times with PBS to assess MHC class I antigen presentation ^20,21^.

#### OT-I CD8 T cell isolation and proliferation activation

CD8^+^ T cells were isolated from the spleens of OT-I transgenic mice using the CD8a^+^ T Cell Isolation Kit (Miltenyi Biotec) according to the manufacturer’s instructions. Purity was confirmed by flow cytometry, and only preparations containing >90% CD3^+^CD8^+^ T cells were used for downstream experiments. In vitro, TPMs from control and RMC-treated mice were incubated in B16-conditioned medium for. After three washes in 1× PBS to remove residual compound, TPMs were cocultured with CFSE-labeled OT-I CD8^+^ T cells (5.0 × 10^6^ cells/mL) for 48 h. T cell activation was defined by the CD44^+^CD62L^−^ phenotype.

### RAW 264.7 transfection

RAW 264.7 macrophages were transiently transfected with pcDNA-DEST40 (Thermo Fisher Scientific) or pD40-His/V5-c-Myc (Addgene plasmid #45597) using Lipofectamine™ 3000 (Thermo Fisher Scientific) according to the manufacturer’s instructions. Briefly, cells were seeded to reach ∼70–80% confluency at the time of transfection, and plasmid DNA was complexed with Lipofectamine 3000 reagent in Opti-MEM. Cells were harvested 48 h post-transfection for downstream analyses.

### Cell viability assay (CCK-8)

Cell viability was assessed using the Cell Counting Kit-8 (MedChemExpress, #HY-K0301) according to the manufacturer’s instructions. Briefly, cells were seeded into 96-well plates and treated as indicated. At the indicated time points, CCK-8 reagent was added to each well and incubated at 37 °C, after which absorbance was measured at 450 nm using a microplate reader.

### Immunofluorescence

For tissue staining, melanoma cryosections were blocked with 1% bovine serum albumin (BSA) and 0.1% Triton X-100 for 1 hour at room temperature. Sections were then incubated overnight at 4 °C with a primary antibody cocktail containing anti-MHC-I (Cell Signaling Technology) and anti-F4/80 (HistoSure). After washing, sections were incubated with appropriate secondary antibodies for 2 hours at 4 °C. Nuclei were counterstained with ProLong™ Gold Antifade Mountant with DAPI (5 μL per section) for 10 minutes. Fluorescent images were acquired using an Axio Scan.Z1 slide scanner (Zeiss).

For cell staining, macrophages isolated from melanoma tissues were seeded on chamber slides (3,000 cells/well; Thermo Fisher). After fixation and blocking, cells were incubated overnight at 4 °C with primary antibodies against c-Myc (Proteintech) and PPIA (Proteintech). Following washes, cells were incubated with secondary antibodies for 2 hours at 4 °C, counterstained with ProLong™ Gold Antifade Mountant with DAPI (5 μL per well) for 10 minutes and imaged using a Leica Stellaris 8 confocal microscope (Leica Microsystems).

### Flow Cytometry

Macrophages were enriched and isolated as described previously and then washed three times with FACS buffer (PBS with 2% FBS, 0.1% sodium azide, 1 mM EDTA)28,33. Cells were suspended in FACS staining buffer at a density of 0.5×106 cells/100 μL and incubated with Fc block (Biolegend) for 10 min on ice. Then cells were stained at 4ºC for 1 hr with anti-mouse antibodies: Pacific Blue-CD45, APC-CD8, PE/cy7-CD11b,, Alexa Fluor-700-F4/80, PE/cy7-CD206,APC-H2Kb(MHC-I), BV785-CD86, BV605-CD44, BV711-CD62L, PE-CD3, BV421-LY6C, FITC-CD163, OVA-ALEX647, PE-SIINFEKL/H-2Kb, BV421-Granzyme B, PE/Dazzle 594-Perforin,, these antibodies are purchased from Biolegend.. After washing three times with FACS buffer, cells were measured or sorted by flow cytometry performed with a Sony ID7000 instrument or a BDAria III Cell Sorter, and data were analyzed with FlowJo version 10.4 (FlowJo, LLC, Ashland, OR).

### Cycloheximide (CHX) Chase and MG132 Assay

Protein stability was analyzed as previously described ^22,23^. Macrophages were treated with RMC-6236 or vehicle for 8 h, followed by cycloheximide (CHX; 50 μg/mL; Selleck) to inhibit protein synthesis. For proteasome inhibition, cells were either pretreated with MG132 (5 μM; Selleck) for 30 min before CHX addition or exposed to MG132 alone for indicated durations to examine c-MYC accumulation kinetics. At each time point, cells were lysed for immunoblotting with anti-MYC antibody. Band intensities were quantified using ImageJ, normalized to time 0, and fitted in GraphPad Prism (v9.0) using a one-phase exponential decay model. Differences in degradation or accumulation kinetics were compared using the extra sum-of-squares F test (p < 0.05).

### AlphaFold3 Structure Prediction and Interface Analysis

The potential interaction between PPIA and MYC was predicted using AlphaFold 3 (https://alphafoldserver.com/). The amino acid sequences of human and mouse PPIA and MYC were obtained from UniProt (P62937 and P01106 for human; P17742 and P01108 for mouse). Default parameters were applied for complex prediction, and model confidence was assessed by the ipTM + pTM composite score. Predicted PPIA–MYC complexes were visualized and analyzed using PyMOL (v2.5, Schrödinger LLC) to identify potential interfacial residues and overlapping regions corresponding to previously reported RMC-binding sites on PPIA.

### RNA-seq and Functional Enrichment Analysis

RNA-seq data from RMC-6236-or vehicle-treated mouse macrophages were analyzed by GSEA (v4.3.3, Broad Institute) using the mouse Hallmark and GO Molecular Function (GOMF) gene sets from MSigDB. Enrichment scores and FDR values were calculated with 1,000 permutations. MYC target genes were further analyzed in STRING (v12.0) using MCL clustering with an inflation parameter of 3 to identify functional gene clusters.

### Chromatin immunoprecipitation followed by sequencing (ChIP–seq)

was performed using the SimpleChIP® Enzymatic Chromatin IP Kit (Magnetic Beads; CST #9003) according to the manufacturer’s instructions. ChIP was performed using antibodies against histone H4 acetylation (H4ac), with matched input controls. An IgG ChIP–seq sample was included as a negative control. Library preparation and sequencing were performed by BGI Genomics. Sequencing reads were quality filtered, trimmed, and aligned to the mouse reference genome (mm10). ChIP–seq data were processed using the nf-core/chipseq pipeline with standard settings. Detailed software versions are listed in **Supplementary Table 2**. Duplicate reads and low-quality alignments were removed prior to downstream analyses. H4ac peaks were identified using MACS3 with input controls. Given the broad distribution of histone acetylation marks, broad peak calling mode was applied. Genome-wide signal tracks were generated and normalized to library size for visualization. For gene-level analyses, promoter regions were defined as ±1 kb around transcription start sites (TSSs). H4ac signal intensity within promoter regions was quantified and compared between experimental groups. Functional annotation of H4ac-associated genes was performed using Gene Ontology (GO) analyses. Integrative Genomic Viewer (IGV ver2.15.2) was used for gene locus visualization.

### ChIP–qPCR (ChIP-PCR)

ChIP samples was prepared using SimpleChIP Enzymatic Chromatin IP Kit (CST #9003) according to the manufacturer‘s protocol. For each sample, 2% of chromatin was reserved as input and processed in parallel. qPCR was performed using SYBR Green chemistry (BIORAD; #1725121) on QuantStudio 3 instrument (Thermofisher). Primer sequences are provided in **Supplementary Table 3**. ChIP enrichment was calculated as percentage of input (%Input).

### IncuCyte imaging analysis

Macrophage phagocytosis and antigen uptake were assessed by live-cell imaging using the Incucyte system (Sartorius). Tumor cells were labeled with CFSE (5 μM, 30 min, 37 °C), washed, and co-cultured with macrophages at a 5:1 effector-to-target ratio. Antigen uptake was measured by incubating macrophages with fluorescent ovalbumin (OVA) from B16-OVA–conditioned supernatant. Phase-contrast and fluorescence images were acquired every 20 min using a 20× objective for phagocytosis assays and a 10× objective for antigen uptake assays. Phagocytosis and antigen uptake were quantified as macrophage-associated intracellular fluorescence using Incucyte software, excluding non-internalized cells by size and intensity thresholds. Experiments were repeated independently at least three times.

### Immunoprecipitation and Western Blotting

Macrophages were purified using anti-mouse F4/80 MicroBeads (Miltenyi Biotec) according to the manufacturer’s protocol. Enriched cells were lysed in RIPA buffer supplemented with Halt™ Protease and Phosphatase Inhibitor Cocktail (100×; Thermo Fisher). Protein concentrations were determined using the Pierce BCA Protein Assay Kit (Thermo Fisher). For immunoprecipitation, whole-cell lysates were prepared from treated cells and processed using the Capturem™ IP & Co-IP Kit (Takara, USA) with an antibody (Novus Biologicals), following the manufacturer’s instructions. Equal amounts of protein (10 μg per lane) were resolved by SDS–PAGE and transferred to polyvinylidene difluoride (PVDF) membranes ^24^. Membranes were blocked in LI-COR blocking buffer for 1 h at room temperature and then incubated with the following primary antibodies (1:1,000 dilution). For IP: Ppia (Proteintech 10720-1-AP), cMyc (Proteintech, #67447-1-Ig) and Mouse (G3A1) IgG1 Isotype Control (CST. #5415). For WB, NLRC5 (E1E9Y) (CST. #72379), c-Myc (CST. #9402), HDAC2 (D6S5P) (CST. #57156), Acetyl-Histone H4 (Lys16) (E2B8W) (CST. #13534), Histone H4 (D2X4V) (CST. #13919), P-ERK (THR202/Tyr204) (CST. #9101), p-S6 (Ser240/244) (CST. #5364), ERK (CST. #4695), S6 (CST. #2217), Vinculin (Santa Cruz #Sc-73614), and β-actin (Sigma. #A3854). IRDye® secondary antibodies (LI-COR Biosciences) were used at 1:10,000 dilution. Signal was detected using a LI-COR Odyssey Fc Imaging System, and band intensities were quantified using Image Studio Lite software (v5.2).

### Quantification and Statistical Analysis

All data are presented as mean ± SEM or mean ± SD, as indicated in the figure legends. GraphPad Prism 9.0 software (GraphPad Software, San Diego, CA, USA) was used to analyze and present the data. Sample size was estimated based on previous experience, sample availability, and previous reported studies. No data were excluded from the data analysis. Normal distribution of data was determined by the Shapiro-Wilk normality test. For statistical comparisons, Unpaired two-tailed t-tests were used to compare two groups; two-way ANOVA followed by Sidak’s multiple comparisons test was used to compare more than two groups. p< 0.05 was considered significant. Numbers per group in the figure legends refer to the number of mice per group.

## Results

### RMC-6236 reinvigorates tumor-associated macrophages and enhances anti-tumor immunity

In the B16-OVA melanoma model, treatment with the pan-RAS inhibitor RMC-6236 resulted in a significant reduction in tumor volume compared with vehicle-treated controls (**Fig. 1A)**, beginning at day 12 and persisting through day 20 (**Fig. 1B**; p < 0.05; **S1A**). In parallel, we examined the phenotypic profiles of circulating and tumor-infiltrating monocytes. RMC-6236 treatment resulted in a reduction of pro-tumoral CD163^+^ monocytes (**Fig. 1C**) and a concomitant upregulation of the co-stimulatory molecules CD80 (**Fig. S1B**) and CD86 (Fig. **1D**) in monocytes, indicating that RMC-6236 reinvigorates macrophage immunocompetence within the tumor microenvironment. We also observed that MHC-I (H-2K^b^) expression was significantly upregulated in monocytes (**Fig. 1E**, p < 0.01), whereas MHC-II (I-A/I-E) remained unchanged (**Fig. S1C**). Consistent with previous reports, the number of CD8^+^ effector T cells also increased following RMC-6236 treatment (**Fig. S1D**).

**Fig. 1.**
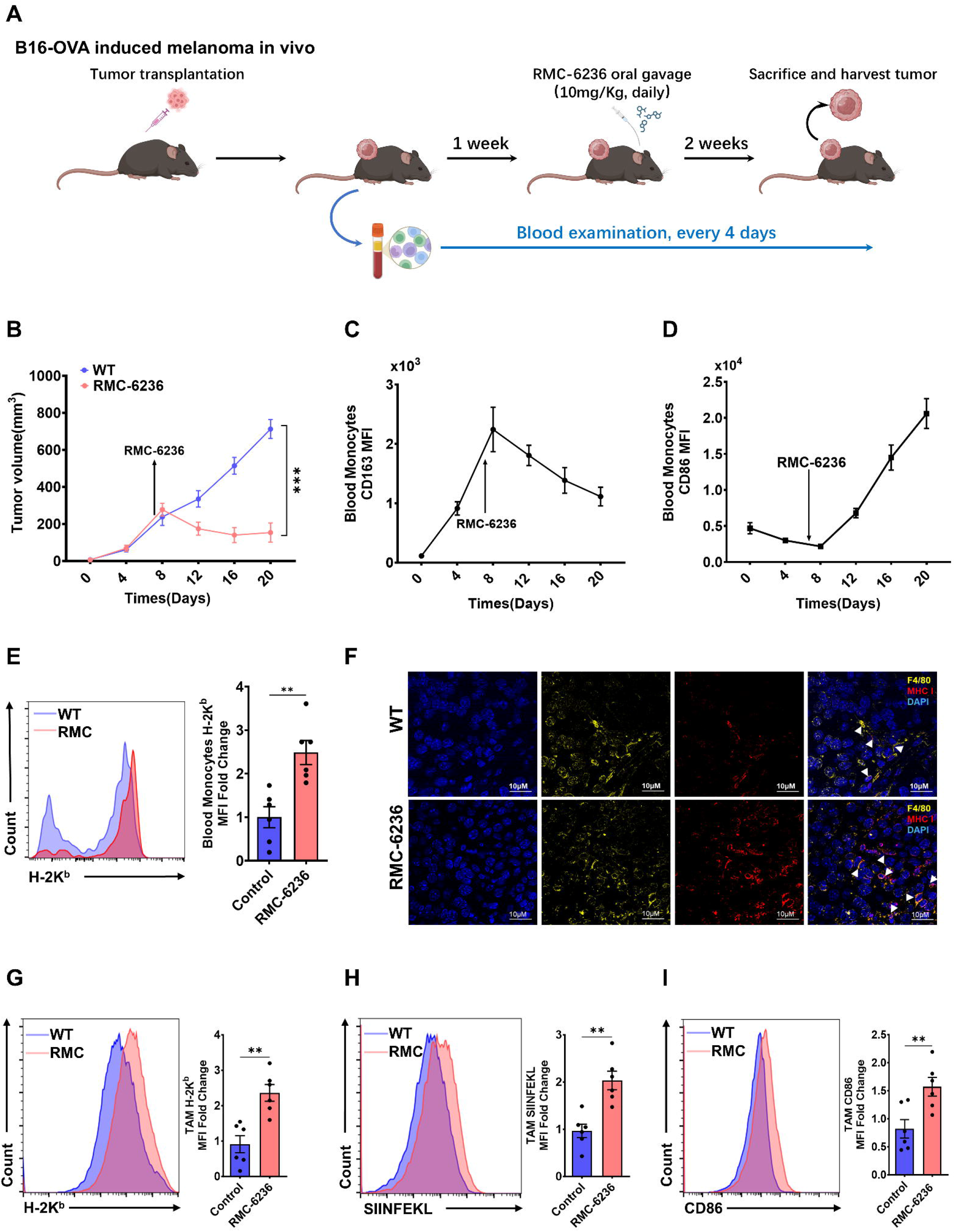
Pan-RAS inhibition reinvigorates macrophage antitumor functions in a B16-OVA melanoma model. **A)** Workflow of B16-OVA–induced melanoma establishment and treatment; **B)** Tumor growth curves of B16 melanoma in vehicle- and RMC-6236-treated mice; **C, D)** Flow cytometric analysis of CD163 **(C)** and CD86 **(D)** expression in monocytes isolated from melanoma tissues; **E)** Endpoint analysis of MHC-I (H-2K^b^) expression in monocytes; **F)** Immunofluorescence analysis of melanoma tissues; scale bars, 10 μm. **G–I)** Tumor-associated macrophage expression of MHC-I **(G)**, antigen-processing capacity (SIINFEKL) **(H)**, and CD86 **(I)** in melanoma tissues from vehicle and RMC-6236-treated mice. Each group included both male and female mice (n = 6). Data are shown as mean ± SEM. *p < 0.05, **p< 0.01, ***p < 0.001 and ****p < 0.0001 using unpaired two-tailed t-tests and Two-way ANOVA followed by Sidak’s multiple comparisons test.

To further assess macrophage responses, we analyzed intratumoral macrophages after tumor collection. RMC-6236 treatment enhanced MHC-I expression (**Fig. 1F, G**) and enhanced OVA-specific antigen presentation, as measured by surface H-2K^b^–SIINFEKL complexes (**Fig. 1H**) in TAMs, without altering total macrophage abundance (**Fig. S1E**). Moreover, RMC-6236 promoted a phenotypic shift from CD206^+^ (M2-like) to CD86^+^ (M1-like) TAMs (**Fig. S1F, Fig. 1I**). As expected, intratumoral CD8^+^ T-cell infiltration was further increased (**Fig. S1G**) ^12^. Notably, RMC-6236 exerted minimal tumor-intrinsic inhibitory effects on B16 melanoma cells in vitro across the tested concentration range (**Fig. S1H**). Pan-RAS inhibition is typically attributed to blockade of RAS–ERK signaling ^8,13^. However, in the B16 model, the antitumor response could not be accounted for by ERK suppression, pointing to an alternative, immune-mediated mechanism (**Fig. S1I**). Importantly, the same in vitro assay demonstrated that RMC-6236 did not impair macrophage viability (Fig. S1H), excluding macrophage toxicity as a contributor to its in vivo effects. Consistently, macrophage functional enhancement was also ERK-independent (**Fig. S1J**).

### RMC-6236 enhances macrophage anti-tumor immunity through NLRC5 activation

To further determine how RMC-6236 augments macrophage anti-tumor function, we first quantified phagocytic capacity in co-cultures of macrophages with CFSE-labelled B16 melanoma cells. We found that RMC-6236 increased macrophage-mediated tumor-cell engulfment compared with vehicle (p<0.01, **Fig. 2A; Fig. S2A**). In a complementary antigen-processing assay using OVA-labelled cargo, RMC-treated macrophages displayed increased antigen processing capacity (p<0.001, **Fig. 2B; Fig. S2B**). Bulk RNA-sequencing of intratumoral macrophages revealed a transcriptional program marked by robust induction of *Nlrc5*, the master regulator of MHC-I antigen presentation ^25^, along with increased *H2-K1* expression (**Fig. 2C,D**). Concordantly, NLRC5 protein expression was elevated both in vitro and in tumor-associated macrophages in vivo following RMC-6236 treatment (**Fig. 2E,F**). In a tumor-conditioned thioglycolate-elicited peritoneal macrophage (TPM) model, RMC-6236 similarly increased CD86 expression, MHC-I surface abundance, and antigen-processing efficiency (**Fig. 2G–I**), while leaving MHC-II unchanged (**Fig. S2C**. Moreover, in TPM–CD8 T-cell co-cultures performed in tumor-conditioned medium (**Fig. 2J**), RMC-6236 enhanced CD8^+^ T-cell activation (**Fig. 2K**) and increased anti-tumor effector molecule production, including TNF-α, Perforin, and Granzyme B (**Fig. 2L–N**).

**Fig. 2.**
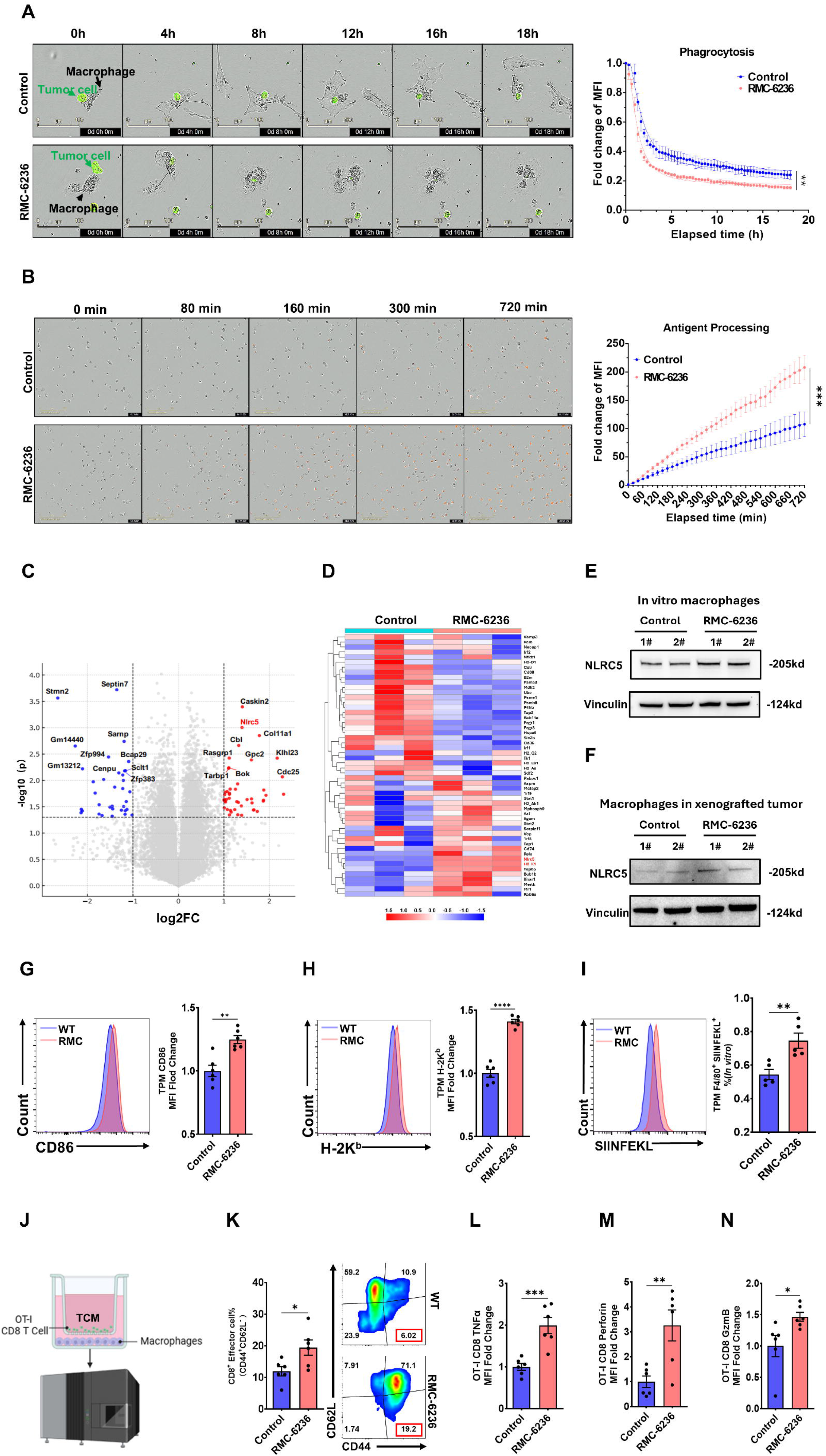
RMC-6236 activates NLRC5 to reawaken macrophage immunity. **A,B)** Incucyte time-lapse imaging of macrophage phagocytosis of CFSE-labelled B16 cells **(A)** and uptake/processing of OVA–Alexa Fluor 647 **(B)**, with quantification. Data are shown as mean ± SD. **p< 0.01 and ***p < 0.001 using unpaired t test (n=3); **C)** RNA-seq volcano plot highlighting Nlrc5 induction by RMC-6236**; D)** Heatmap of differentially expressed antigen-presentation/immune genes**; E,F)** Immunoblot of NLRC5 in macrophages treated in vitro **(E)** and intratumoral macrophages **(F); G–I)** Flow cytometry of intratumoral macrophages CD86 **(G)**, MHC-I **(H)**, and SIINFEKL presentation **(I); J)** Schematic of TPM– OT-I CD8 T-cell co-culture in tumor-conditioned medium**; K)** CD8 T-cell activation (CD44/CD62L)**; L–N)** CD8^+^ T-cell cytotoxic effector molecules, including TNF-α**(L)**, perforin**(M)** and granzyme B**(N)**. Data are shown as mean ± SEM. *p < 0.05, **p< 0.01, ***p < 0.001 and ****p < 0.0001 using unpaired two-tailed t-tests (n≥3) and Two-way ANOVA followed by Sidak’s multiple comparisons test.

### RMC-6236–induced anti-tumor immunity is macrophage-dependent

Previous studies showed that pan-RAS inhibition can enhance CD8^+^ T-cell function^11,12^. To determine whether RMC-6236 acts directly on T cells or relies on macrophage activation, we implanted B16-OVA melanoma cells into T-cell-deficient nude mice (**Fig. 3A**). Remarkably, RMC-6236 retained tumor-suppressive activity in the absence of T cells, with RMC-treated nude mice showing reduced tumor size compared with vehicle (**Fig. 3B,C**). Tumor-associated macrophages underwent a phenotypic shift from CD206^+^ to CD86^+^ states (**Fig. 3D,E**) with increased MHC-I expression and antigen-processing activity, while MHC-II remained unchanged (**Fig. 3F-I**). To directly test macrophage dependence, we depleted monocytes/macrophages using anti-CSF1R (**Fig. 3J**). Monocyte abundance was markedly reduced at endpoint. (p<0.0001, **Fig. S2D**). Under these conditions, the anti-tumor effect of RMC-6236 was abolished (**Fig. 3K**), and CD8^+^ T-cell abundance and cytotoxic function were no longer increased (**Fig. 3L–O**), indicating that RMC-6236 requires monocytes/macrophages, rather than T cells, to elicit its immune-enhancing effects ^12^.

**Fig. 3.**
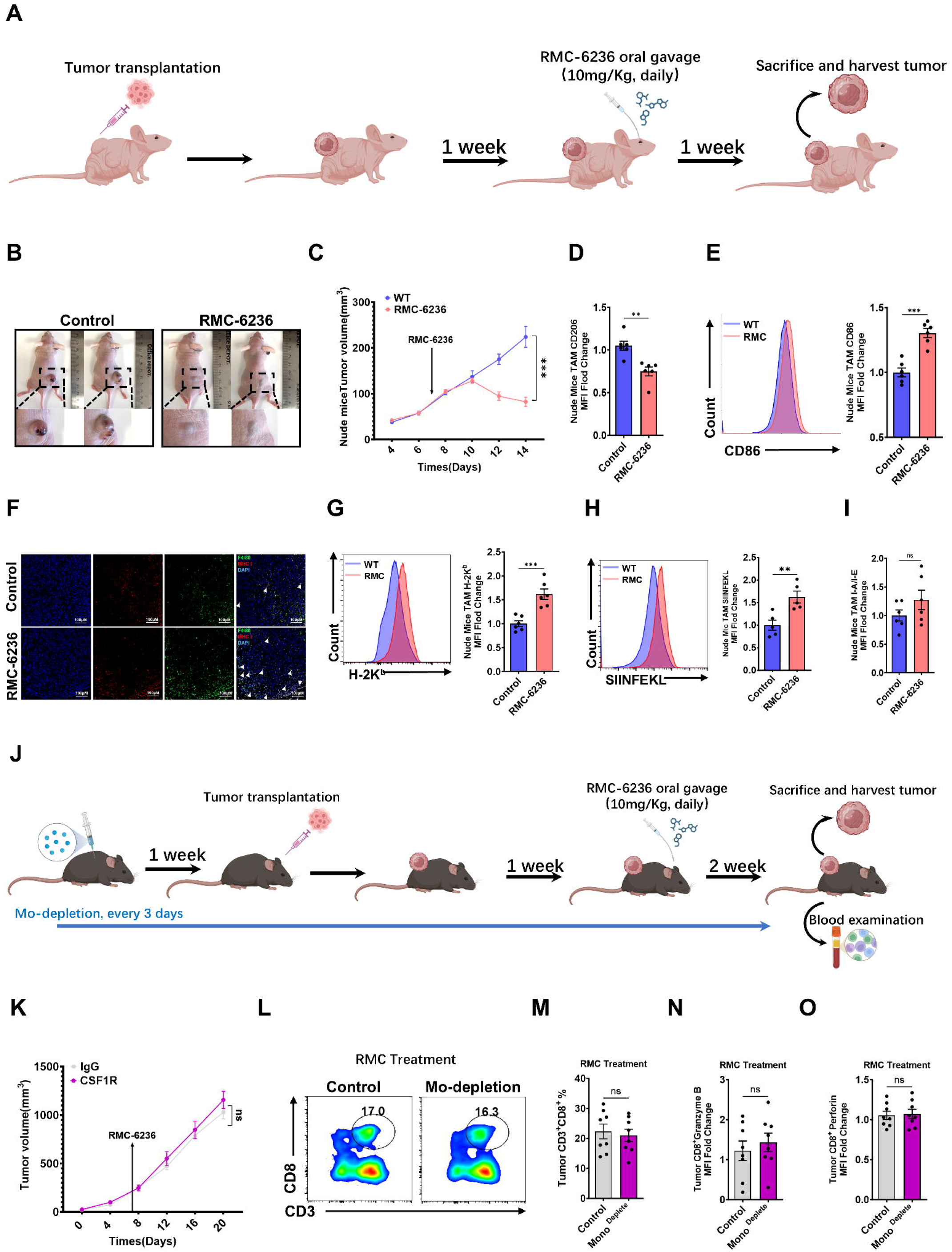
RMC-6236 suppresses tumors through monocytes/macrophages independent of T cells. **A)** Experimental scheme for B16-OVA transplantation in nude mice followed by RMC-6236 oral gavage (10 mg/kg, daily)**; B) C**omparison of tumor size between vehicle and RMC-6236–treated nude mice**; C)** Tumor growth curves**; D,E)** Flow-cytometry analysis of TAM CD206 **(D)** and CD86 **(E)** with vehicle or RMC-6236**; F)** Immunofluorescence showing macrophages (F4/80, green) and MHC-I (red) with DAPI (blue)**; G–I)** Flow-cytometry quantification in TAMs of MHC-I (H-2K^b^) **(G)**, OVA-specific presentation (SIINFEKL) **(H)**, and MHC-II (I-A/I-E) **(I); J)** Schematic of anti-CSF1R– mediated monocyte/macrophage depletion during RMC-6236 treatment; **K)** Tumor growth with RMC-6236 plus IgG or anti-CSF1R**; L, M)** Intratumoral CD8^+^ T-cell frequency after monocyte depletion**; N,O)** Granzyme B **(N)** and perforin **(O)** in tumor CD8^+^ T-cells. Data are shown as mean ± SEM. *p < 0.05, **p< 0.01, ***p < 0.001 and ****p < 0.0001 using unpaired two-tailed t-tests and Two-way ANOVA followed by Sidak’s multiple comparisons test (n≥6).

### RMC-6236 epigenetically activates Nlrc5 in macrophages by suppressing Myc signaling

To define how RMC-6236 induces NLRC5 and enhances antigen presentation, we performed GSEA on macrophage RNA-seq from vehicle vs RMC-6236-treated mice. Myc target genes were significantly enriched in controls **(Fig. 4A; Fig.S2A**). Consistently, RMC-6236 reduced Myc protein in macrophages in vitro and in xenograft tumors (**Fig.S2B**), and downregulated multiple Myc target transcripts (**Fig. 4B)**.

**Fig. 4.**
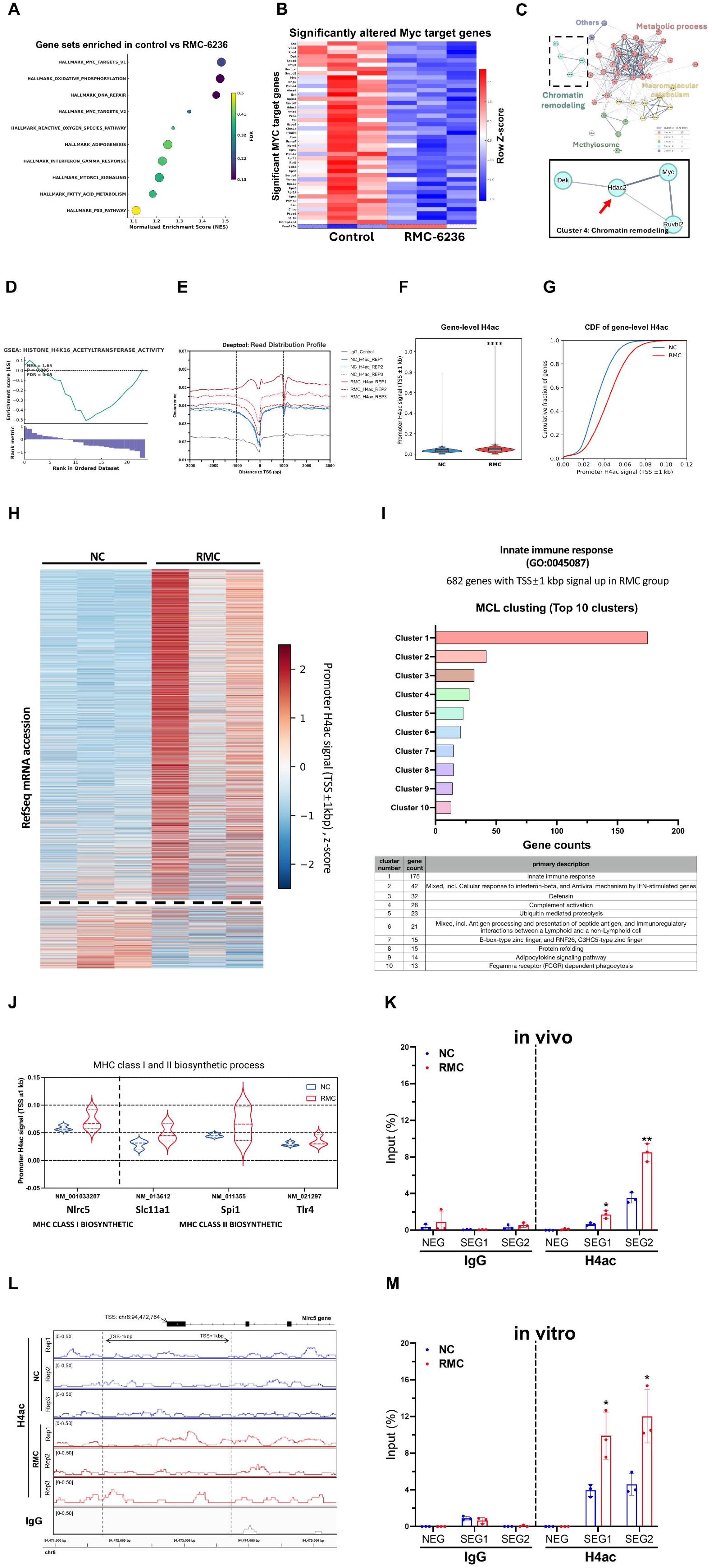
RMC-6236 suppresses Myc programs and increases promoter H4 acetylation in macrophages. **A)** GSEA of RNA-seq from macrophages isolated from vehicle-or RMC-6236–treated mice; each dot denotes a gene set (size, leading-edge genes; color, FDR); **B)** Heatmap of significantly altered Myc target genes (row Z-score) in vehicle vs RMC-6236 macrophages; **C)** STRING protein–protein interaction network and clustering of altered Myc targets, highlighting a chromatin-remodeling module containing Hdac2; **D)** GSEA showing enrichment of the GO term histone H4 acetyltransferase activity in RMC-6236 vs vehicle; **E)** Metagene profile of H4ac ChIP–seq signal across genes aligned to TSSs in primary mouse macrophages treated ex vivo with vehicle (NC) or RMC, with IgG control; **F)** Violin plots of promoter H4ac signal (TSS ±1 kb) in NC vs RMC. Center lines indicate medians; boxes denote interquartile ranges. ****p < 0.0001 using a two-sided Mann– Whitney U test; **G)** CDF of promoter H4ac signal (TSS ±1 kb) showing a rightward shift with RMC. Statistical significance was assessed using a two-sided Mann–Whitney U test; **H)** Heatmap of promoter H4ac signal (TSS ±1 kb) across transcripts ordered by change between conditions; **I)** Top ten MCL clusters from RMC-upregulated innate immune response genes (GO:0045087); **J)** Promoter H4ac signal (TSS ±1 kb) for selected MHC-related genes (including Nlrc5); **K)** IGV tracks of H4ac signal at the Nlrc5 locus in NC vs RMC primary macrophages (three biological replicates per condition; same y-axis scale; IgG control). **L, M**) ChIP–qPCR validation of increased H4ac at Nlrc5 regulatory regions defined by ChIP–seq (**Fig. S3F**) in vehicle or RMC-6236–treated macrophages, including primary macrophages treated ex vivo (n = 3 per group). Data are shown as mean ± SD. *p < 0.05 and **p< 0.01 using unpaired two-tailed t-tests.

Following STRING analysis of Myc-responsive genes, highlighted clusters linked to metabolism and chromatin regulation (**Fig. 4C; Fig. S2C**). Given robust *Nlrc5* induction, we focused on the chromatin remodeling, Notably, Hdac2 (histone deacetylase 2), a key repressor of chromatin accessibility, was significantly downregulated by RMC-6236 (**Fig. 4B,C**), and GSEA showed enrichment of “Histone H4K16 acetyltransferase activity” in RMC-treated macrophages (**Fig. 4D**). Because HDAC2 restrains H4 acetylation, its reduction would be expected to enhance acetyl-H4K16-mediated epigenetic activation and facilitate Nlrc5 transcription.

We therefore performed acetyl-H4K16 (H4ac) ChIP-seq in vehicle and RMC-treated mice macrophages. PCA showed clear separation between groups (**Fig. S2D**). RMC-6236 increased H4ac around transcription start sites (TSSs) (**Fig. 4E; Fig. S2E**) with a global increase in promoter-associated H4ac (±1 kb of TSS) by violin and cumulative distribution function (CDF) analyses **(Fig. 4F–G**). Heatmap analysis revealed that transcripts exhibiting increased promoter-associated H4ac upon RMC treatment markedly outnumbered those with decreased signals, suggesting a global bias toward promoter hyperacetylation (**Fig. 4H**). Among genes with increased promoter H4ac, 682 innate immune response genes were identified and clustered (MCL, inflation=2), revealing modules enriched for macrophage activation and effector functions (**Fig. 4I**). Notably, promoter-associated H4ac was elevated at genes involved in MHC biosynthetic process, including *Nlrc5*, a well-established regulator of MHC class I ^25^, as well as *Slc11a1, Spi1*, and *Tlr4*, which have been linked to MHC-II pathway (**Fig. 4J**). Consistent with our cellular data showing selective induction of MHC class I but not class II (**Fig. S1C, 2C**). Genome browser visualization confirmed increased H4ac enrichment at the *Nlrc5* promoter across replicates **(Fig. 4K**). ChIP–qPCR was performed on macrophages treated in vitro with vehicle or RMC-6236. Based on IGV inspection of the *Nlrc5* locus, we identified two promoter-proximal H4ac-enriched regions (SEG1 and SEG2) and a no-peak control region (NEG) (**Fig. S2F**). RMC-6236 significantly increased H4ac enrichment at SEG1 and SEG2 compared with vehicle, validating the ChIP–seq results (**Fig. 4L**). This increase was recapitulated in primary murine macrophages treated ex vivo with RMC-6236, supporting a cell-intrinsic gain of H4ac occupancy at Nlrc5 (**Fig. 4M**).

### RMC-6236 promotes Myc degradation to unleash antigen presentation in macrophages

We next tested causality in the Myc–HDAC2–H4ac pathway. Genetic Myc knockdown using Myc-RIBOTAC reduced MYC and HDAC2 while increasing NLRC5, and enhanced MHC-I antigen presentation; importantly, it blunted the additional effect of RMC-6236 (**Fig. 5AB,**). The Myc inhibitor 10058-F4 phenocopied these effects (**Fig. 5CD,**). Conversely, transient Myc overexpression increased HDAC2, reduced histone H4 acetylation, and suppressed NLRC5 expression and MHC-I presentation, effects that were reversed by HDAC2 inhibition (HDAC2-IN-2) in Myc-overexpressing RAW264.7 macrophages (**Fig. S4AB,**). Consistently, HDAC2 inhibition eliminated the dependence of RMC-6236 on upstream Myc suppression in primary macrophages (**Fig. 5EF,**). Finally, blocking histone acetyltransferase activity with PU139 abolished NLRC5 induction and MHC-I enhancement driven by Myc inhibition and by RMC-6236 (**Fig. 5GH,; Fig. S4C–F**), establishing histone H4 acetylation as the key downstream effector.

**Fig. 5.**
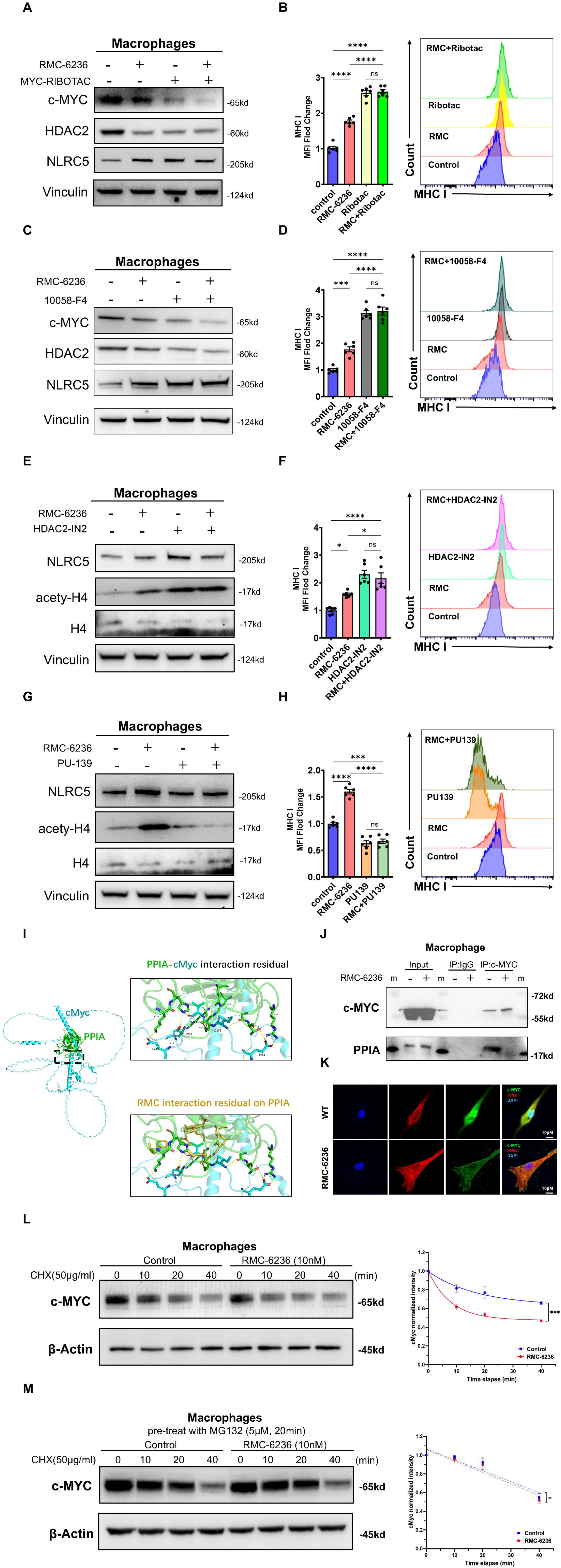
RMC-6236 engages a Myc–HDAC2 axis and perturbs PPIA–Myc interactions in macrophages. **A–H**) Immunoblot of c-MYC, HDAC2 and NLRC5 and flow cytometry analysis of MHC-I in macrophages treated with vehicle or RMC-6236, in the absence or presence of MYC-RIBOTAC (**A,B**), 10058-F4 (**C,D**), HDAC2-IN2 (**E,F**) or PU-139 (**G,H**). **I**) Structural modelling highlighting residue overlap between the PPIA–c-Myc interface and the RMC interaction surface on PPIA; **J**) Co-immunoprecipitation of c-MYC showing associated PPIA in macrophages with or without RMC-6236; **K**) Immunofluorescence of c-MYC and PPIA in macrophages treated with vehicle or RMC-6236; **L**) **Cycloheximide (CHX) chase assay in macrophages treated with vehicle or RMC-6236**.; **M**) CHX chase assay in macrophages pre-treated with MG132. Data are shown as mean ± SD. *p < 0.05, **p< 0.01, ***p < 0.001 and ****p < 0.0001 using extra sum-of-squares F test (n=3).

RMC-6236 is a clinical-stage RAS(ON) inhibitor, yet at 10 nM it did not measurably alter RAS–ERK signaling in macrophages (**Fig. S4G**), suggesting a non-canonical, cell-type– specific mechanism. Because RMC compounds recruit PPIA (Cyclophilin A) as part of their molecular-glue activity [38589574], and given that PPIA is a ubiquitously expressed intracellular immunophilin, we hypothesized that PPIA may stabilize MYC in macrophages. AlphaFold multimer modeling predicted a direct PPIA–MYC interaction in human and mouse, revealing an interface that partially overlaps with residues previously implicated in RMC–PPIA binding (**Fig. 5I; Fig. S4H**). Co-IP and immunofluorescence confirmed that RMC-6236 weakened PPIA–MYC association (**Fig. 5J,K**). Functionally, RMC-6236 shortened MYC half-life in CHX-chase assays (**Fig. 5L**), and proteasome blockade with MG132 caused rapid MYC accumulation and erased the differential degradation kinetics (**Fig. 5M; Fig. S4I**), indicating accelerated proteasomal turnover.

## Discussion

The development of next-generation pan-RAS(ON) inhibitors, such as RMC-6236 and RMC-7977, has overcome longstanding barriers to pharmacologic RAS targeting by enabling inhibition of active RAS across isoforms and genotypes ^8,13^. Beyond their direct anti-tumor efficacy, accumulating preclinical evidence suggests that pan-RAS(ON) inhibition is accompanied by broad remodeling of the tumor immune microenvironment. In particular, treatment of KRAS ^G12D^-driven tumors with pan-RAS inhibitors has been shown to increase intratumoral CD8^+^ T cell infiltration and enhance responsiveness to immune checkpoint blockade^12^, underscoring a previously underappreciated immunomodulatory dimension of RAS-targeted therapy.

Given their central role at the intersection of innate and adaptive immunity within the tumor microenvironment, macrophages are uniquely positioned to translate immunomodulatory signals into effective antitumor immune responses ^26^. Tumor-associated macrophages (TAMs) are highly plastic and can be therapeutically reprogrammed, making them attractive targets for cancer therapy. However, in many tumors TAMs display impaired antigen-presenting capacity, with reduced MHC-I and MHC-II expression ^27,28^. Currently, one prominent strategy is to disrupt the CD47–SIRPα “don’t-eat-me” checkpoint: SIRPα is a myeloid-enriched inhibitory receptor that restrains phagocytosis upon binding CD47 ^29,30^. Blocking this axis with CD47-targeting antibodies releases the phagocytic brake, promoting SIRPα-dependent antibody-dependent cellular phagocytosis (ADCP) and, in some settings, repolarizing TAMs toward an M1-like pro-inflammatory state ^1,31^.

In this study, we identify a distinct macrophage-centered immune mechanism underlying pan-RAS inhibition that operates independently of tumor-intrinsic RAS dependency. In a RAS–wild-type melanoma context, the therapeutic activity of RMC-6236 cannot be fully accounted for by direct tumor cell inhibition or by T cell–intrinsic mechanisms alone. Instead, our findings support a model in which macrophages serve as critical mediators of pan-RAS–induced antitumor immunity through restoration of antigen presentation capacity, thereby reshaping the tumor immune microenvironment. These data suggest that macrophages act as the primary effector and initiating compartment for RMC-6236– induced antitumor immunity, while T cells function as downstream amplifiers rather than obligate mediators.

Mechanistically, our data indicate that pan-RAS inhibition in macrophages engages a noncanonical signaling axis largely uncoupled from classical RAS–ERK blockade. Rather than suppressing ERK activity, RMC-6236 attenuates a MYC–HDAC2–dependent epigenetic program that couples transcriptional output to chromatin accessibility. Consistent with prior studies implicating MYC as a key driver of alternative (M2-like) macrophage activation ^32,33^, our findings extend this paradigm by uncovering a previously underappreciated role for MYC in restraining antigen presentation programs. By relieving MYC–HDAC2–mediated epigenetic constraints, pan-RAS inhibition enhances histone H4K16 acetylation at regulatory elements of the *Nlrc5* locus, thereby promoting NLRC5-driven MHC-I antigen presentation and shifting macrophages toward a more immunostimulatory state. Whether MYC constrains antigen presentation directly at immune gene loci or indirectly through broader metabolic and chromatin remodeling remains an important question for future studies.

Although antigen presentation represents a principal functional output examined in this study, additional macrophage programs—including cytokine production, metabolic remodeling, or complement activation—may also be influenced downstream of MYC suppression. Moreover, MYC is a central oncogenic driver in tumor cells ^34^, raising the intriguing possibility that pan-RAS inhibitors could coordinately modulate MYC-dependent programs across both tumor and immune compartments. Elucidating how pan-RAS inhibition synchronizes MYC signaling across distinct cellular contexts therefore represents an important avenue for future investigation.

An important mechanistic question raised by our findings concerns how pan-RAS inhibition selectively attenuates MYC programs in macrophages. RMC-6236 functions as a molecular glue that stabilizes a ternary complex between active RAS and the immunophilin cyclophilin A (PPIA), thereby constraining RAS signaling output ^8,13^ Our data suggest that PPIA may also interact with MYC, raising the possibility that engagement of PPIA by RMC-6236 perturbs PPIA–MYC interactions, thereby potentially altering MYC stability in macrophages.

As an intracellular immunophilin, PPIA plays broad roles in protein folding, chaperoning, and proteostasis^35^. Accordingly, pharmacologic targeting of PPIA by molecular-glue pan-RAS inhibitors may exert effects beyond canonical RAS signaling, particularly in cellular contexts in which RAS activity is not pathologically hyperactivated. Consistent with prior reports linking PPIA to macrophage phagocytosis and inflammatory programming^36-38^, perturbation of PPIA-centered networks may contribute to the immune reprogramming effects observed with RMC-6236. Taken together, these considerations suggest that engagement of PPIA expands the pharmacologic scope of molecular-glue pan-RAS inhibitors toward previously underappreciated immune-regulatory pathways. Nevertheless, while our data support a role for PPIA in regulating MYC stability, the molecular determinants governing PPIA–MYC regulation and their modulation by RMC-6236 remain to be fully defined. It remains possible that PPIA-dependent effects on MYC stability are indirect and integrated within broader proteostasis networks. Whether this macrophage-intrinsic epigenetic program represents a class effect of molecular-glue pan-RAS(ON) inhibitors, or exhibits compound-specific features, will require comparative studies.

Together, our study broadens the paradigm of pan-RAS inhibition by revealing that the clinically advancing pan-RAS(ON) inhibitor RMC-6236 functions as an epigenetic immunomodulator that restores macrophage antigen presentation through the Myc– HDAC2–NLRC5 axis. This previously unrecognized, macrophage-intrinsic mechanism provides a rationale for leveraging RMC-6236 to reprogram the tumor immune microenvironment across tumor genotypes and highlights macrophage epigenetic plasticity as a tractable target for cancer immunotherapy.

## Author contributions

J.S., and D.-Y.C., conceived and designed the experiments. Z.C., J.S., and D.-Y.C acquired data with assistance from M.-T.C. (tissue sections), S.D.J helped the flow sorting, C.-B.Z., H.-D S.-C., W.-N. L., W.F., Y.Y., (macrophages isolation, cell culture, animal husbandry, flow cytometer, mice genotyping). J.S., and D.-Y.C., designed the CD8 experiments. J.S. processed the RNA and ChIP data on macrophages.. D.-Y.C., J.S., H.-P.Y., Z.K. contributed to manuscript editing.

## Acknowledgements

This study was supported by AHA grants 23CDA1052548 (D.Y.C).

The National Natural Science Foundation of China, Grant No. 32272825 (Z.C), 32472851 (Z.C), 32400582 (J.S).

## Conflict of Interest

The authors declare that there is no conflict of interest regarding the publication of this article.

---

**Schematic: pan-RAS inhibition relieves macrophage suppression and promotes a switch toward phagocytosis and antigen presentation in the tumor microenvironment**.

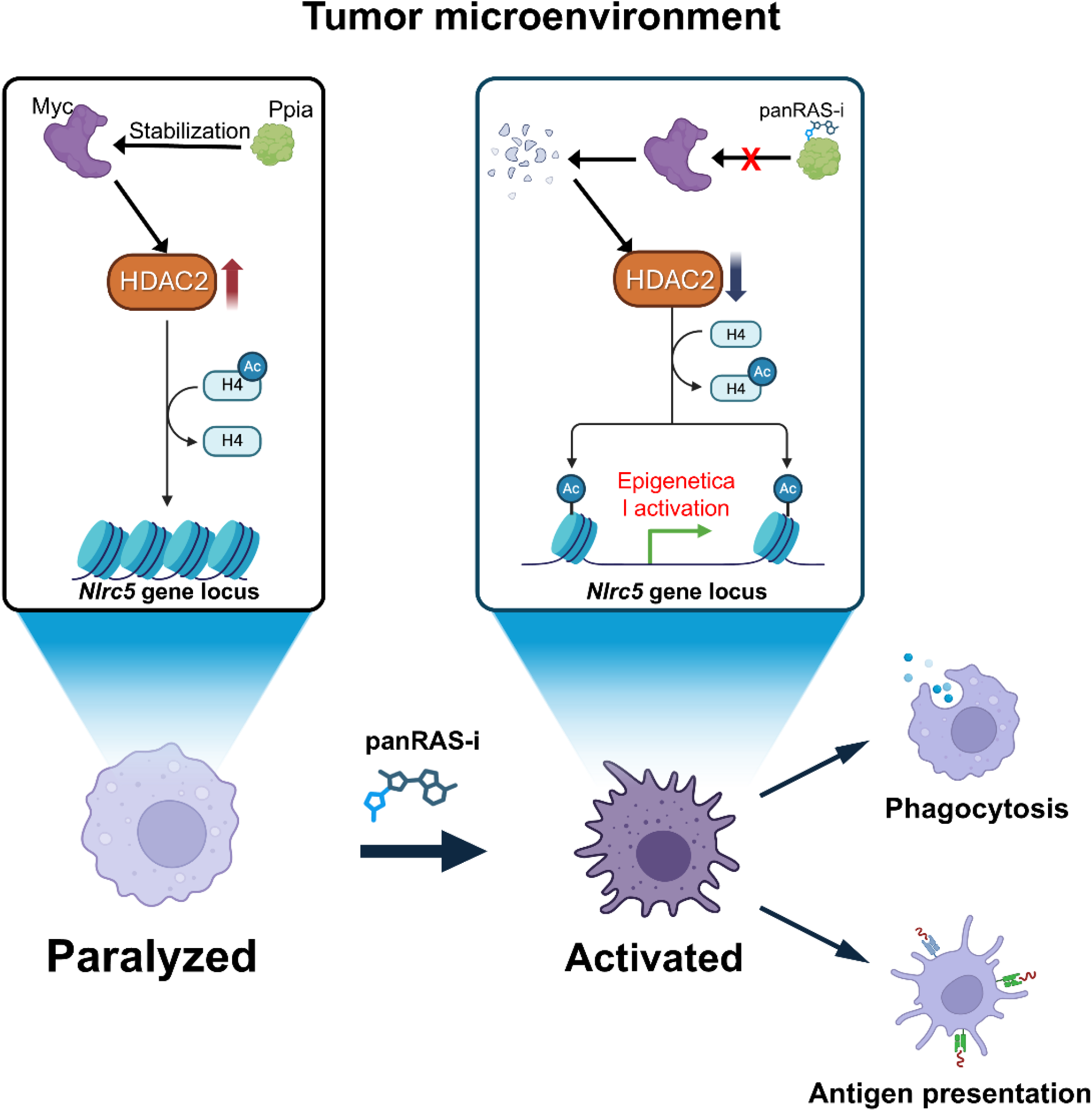

**Supplemental Fig. 1. RMC-6236 activates myeloid cells and enhances CD8 T-cell responses. A)** Representative endpoint B16 tumors from vehicle and RMC-6236–treated mice; **B,C)** Flow cytometry analysis of CD80 **(B)** and MHC-II (I-A/I-E) **(C)** expression in blood monocytes; **D)** Frequency of circulating CD8 effector T cells (CD44^+^CD62L^−^); **E)** Gating strategy and frequency of tumor-infiltrating macrophages (CD11b+F4/80+); **F)** TAM CD206 expression; **G)** Tumor-infiltrating CD8 T cells; **H)** Viability of B16 cells and macrophages treated with increasing doses of RMC-6236. Data are shown as mean ± SD (n=3); **I)** Immunoblot analysis of p-ERK and p-S6 in macrophages following RMC-6236 dose titration; **J)** Flow cytometry analysis of p-ERK in macrophages. Each group included both male and female mice (n = 6). Data are shown as mean ± SEM. ^*^p < 0.05, ^**^p< 0.01, ^***^p < 0.001 and ^****^p < 0.0001 using unpaired two-tailed t-tests and Two-way ANOVA followed by Sidak’s multiple comparisons test.

**Supplementary Fig. 2. RMC-6236 enhances macrophage phagocytosis and antigen processing. A**,**B)** Representative Incucyte videos of macrophage phagocytosis of CFSE-labelled B16 melanoma cells **(A)** and uptake/processing of OVA–Alexa Fluor 647 **(B)** under control or RMC-6236 conditions; **C)** Flow cytometry analysis of MHC-II (I-A/I-E) expression in macrophages cultured with tumor-conditioned medium; **D)** Flow cytometry analysis of circulating CD11b^+^Ly6C^+^ after CSF1R antibody–mediated depletion. Data are shown as mean ± SEM. ^*^p < 0.05, ^**^p< 0.01, ^***^p < 0.001 and ^****^p < 0.0001 using unpaired two-tailed t-tests and Two-way ANOVA followed by Sidak’s multiple comparisons test (n ≥ 5).

**Supplemental Fig. 3. RMC-6236 suppresses MYC signaling and remodels H4ac at the Nlrc5 locus**.

**A)** GSEA enrichment plot for HALLMARK_MYC_TARGETS_V1 comparing control and RMC-6236–treated macrophages; heatmap below shows expression of leading-edge MYC target genes across samples; **B)** Immunoblot analysis of Myc in macrophages treated with vehicle or RMC-6236 in vitro (left) and in vivo (right); **C)** Bubble plots of GO enrichment for each STRING-defined cluster of differentially expressed MYC target genes (dot size, gene count; color, FDR)**; D)** PCA of H4ac ChIP–seq profiles from **primary mouse macrophages treated with vehicle (NC) or RMC** (each dot, one biological replicate)**; E)** Metagene profile of H4ac ChIP–seq signal across annotated genes**; F)** IGV tracks of H4ac ChIP–seq signal at the Nlrc5 locus in NC- and RMC-treated primary macrophages. Two RMC-enriched regions (SEG1 and SEG2) were selected for ChIP–qPCR, with a no-peak region (NEG) as a negative control.

**Supplementary Fig. 4. Genetic and pharmacologic validation of the Myc–HDAC2– H4ac pathway controlling NLRC5 and MHC-I**.

**A–D**) Analysis of c-MYC, HDAC2, NLRC5 and acetyl-H4 and MHC-I quantification in RAW264.7 cells with c-Myc overexpression treated with RMC-6236 (**A,B**) or HDAC2-IN2 (**C,D**); **E–H**) Immunoblot analysis of c-MYC, NLRC5 and acetyl-H4 and MHC-I quantification in macrophages treated with MYC-RIBOTAC (**E,F**) or 10058-F4 (**G,H**) in the absence or presence of PU-139; **I**) Immunoblot of p-ERK and total ERK in macrophages treated with vehicle or RMC-6236; **J**) Predicted human and mouse PPIA–MYC interaction models; **K**) MG132 time-course showing c-MYC accumulation in macrophages treated with vehicle or RMC-6236, with quantification. Data are shown as mean ± SD. ^*^p < 0.05, ^**^p< 0.01, ^***^p < 0.001 and ^****^p < 0.0001; extra sum-of-squares F test (n = 3).

**Supplementary Table 1.** Complete list of inhibitors and Reagents.

| Name | Catalog Number | Vendor |
| --- | --- | --- |
| HDAC2-IN-2 | HY-164050 | MedChemExpress |
| PU139 | HY-124696 | MedChemExpress |
| 10058-F4 | HY-12702 | MedChemExpress |
| MYC-RIBOTAC | HY-153713 | MedChemExpress |
| RMC-6236 | HY-148439 | MedChemExpress |
| Dimethyl sulfoxide (DMSO) | HY-Y0320 | MedChemExpress |
| Cycloheximide (CHX) | S7418 | Selleck |
| MG132 | S2619 | Selleck |
| Cell Counting Kit-8 (CCK-8) | HY-K0301 | MedChemExpress |

**Supplementary Table 2.**
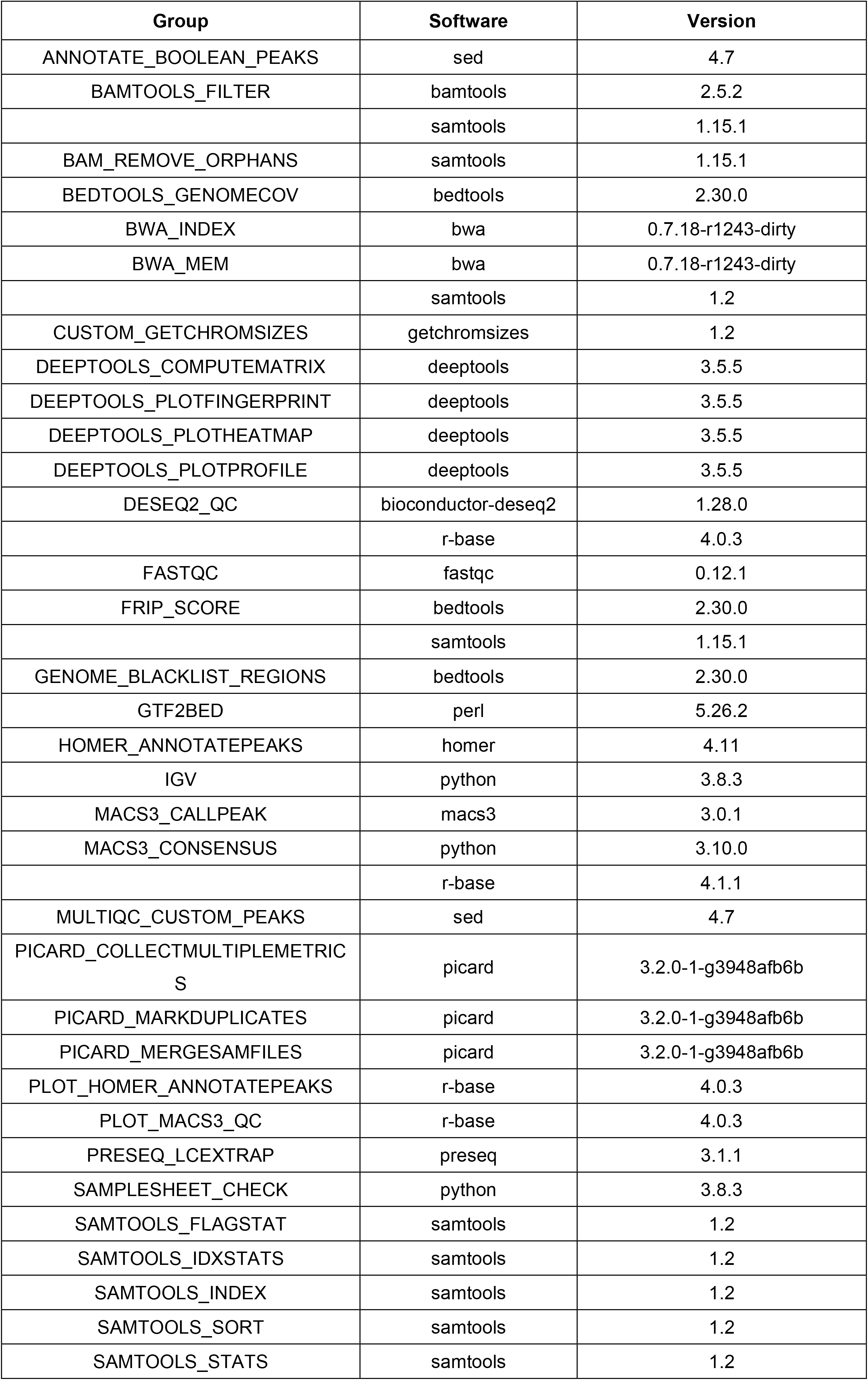
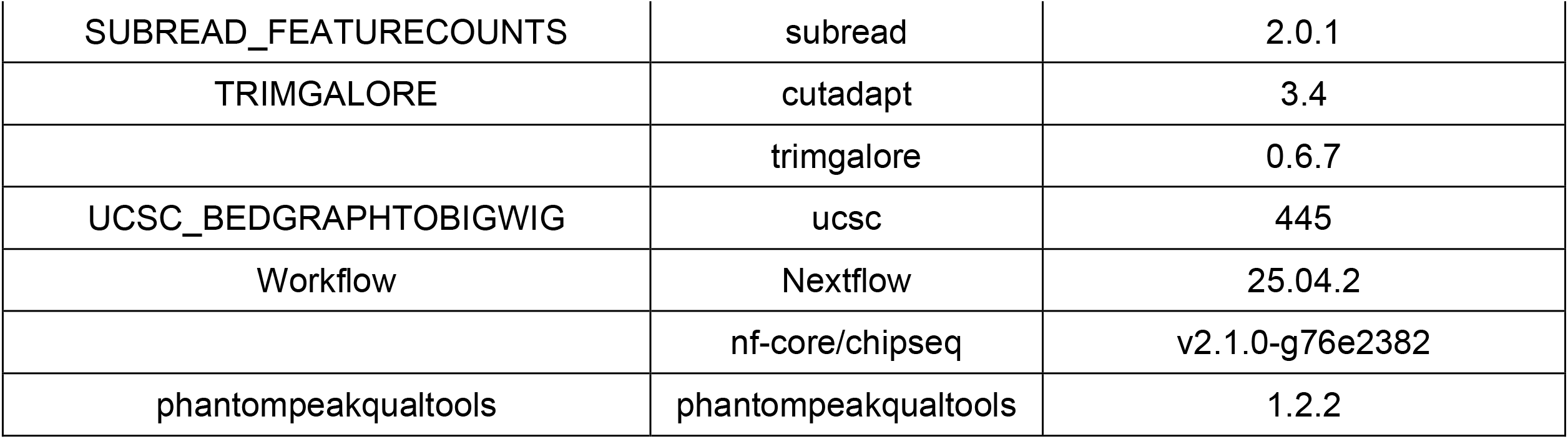
Softwares using the nf-core/chipseq pipeline.

**Supplementary Table 3.**
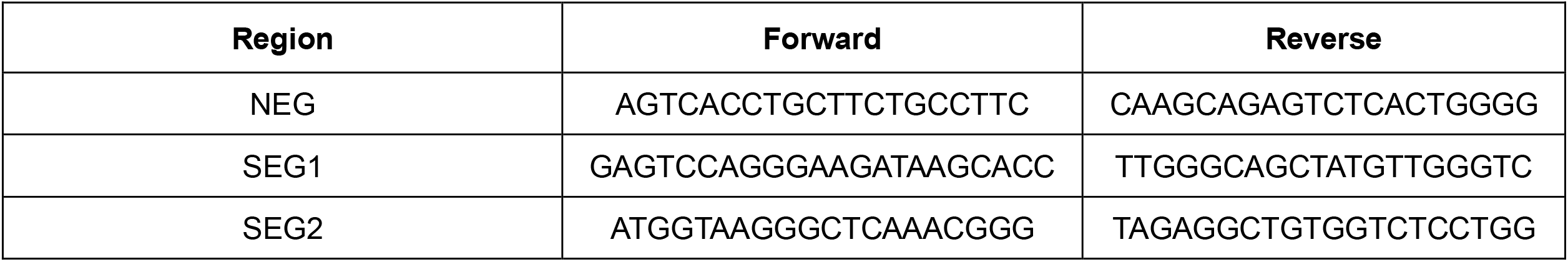
Primer for ChIP-qPCR.

## Notes

### Competing Interest Statement

The authors have declared no competing interest.

