## Supplementary figures and images for "Macrophage Antigen Presentation Is Unleashed by Pan-RAS Inhibition to Promote Antitumor Immunity"

### Supplemental figures

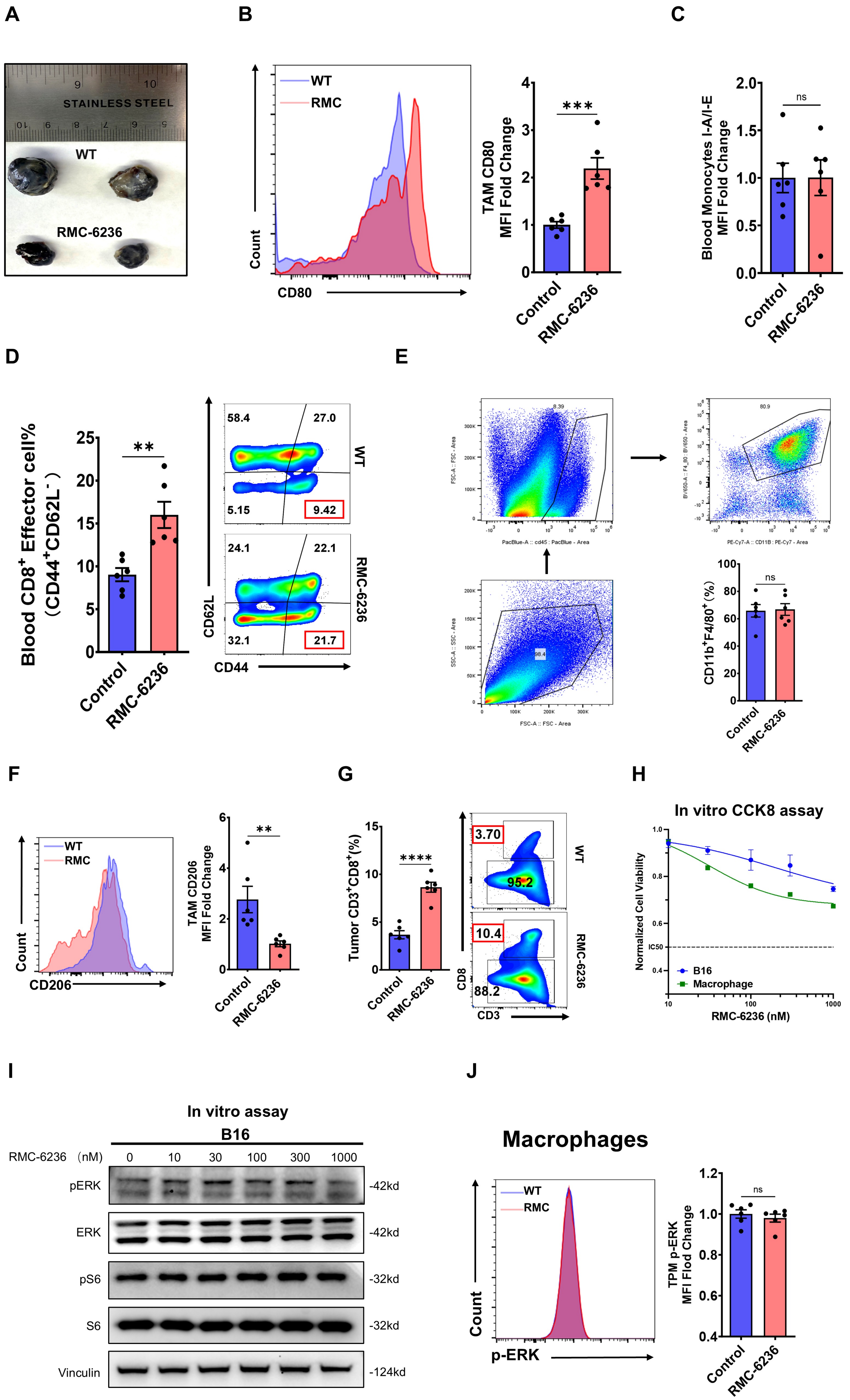

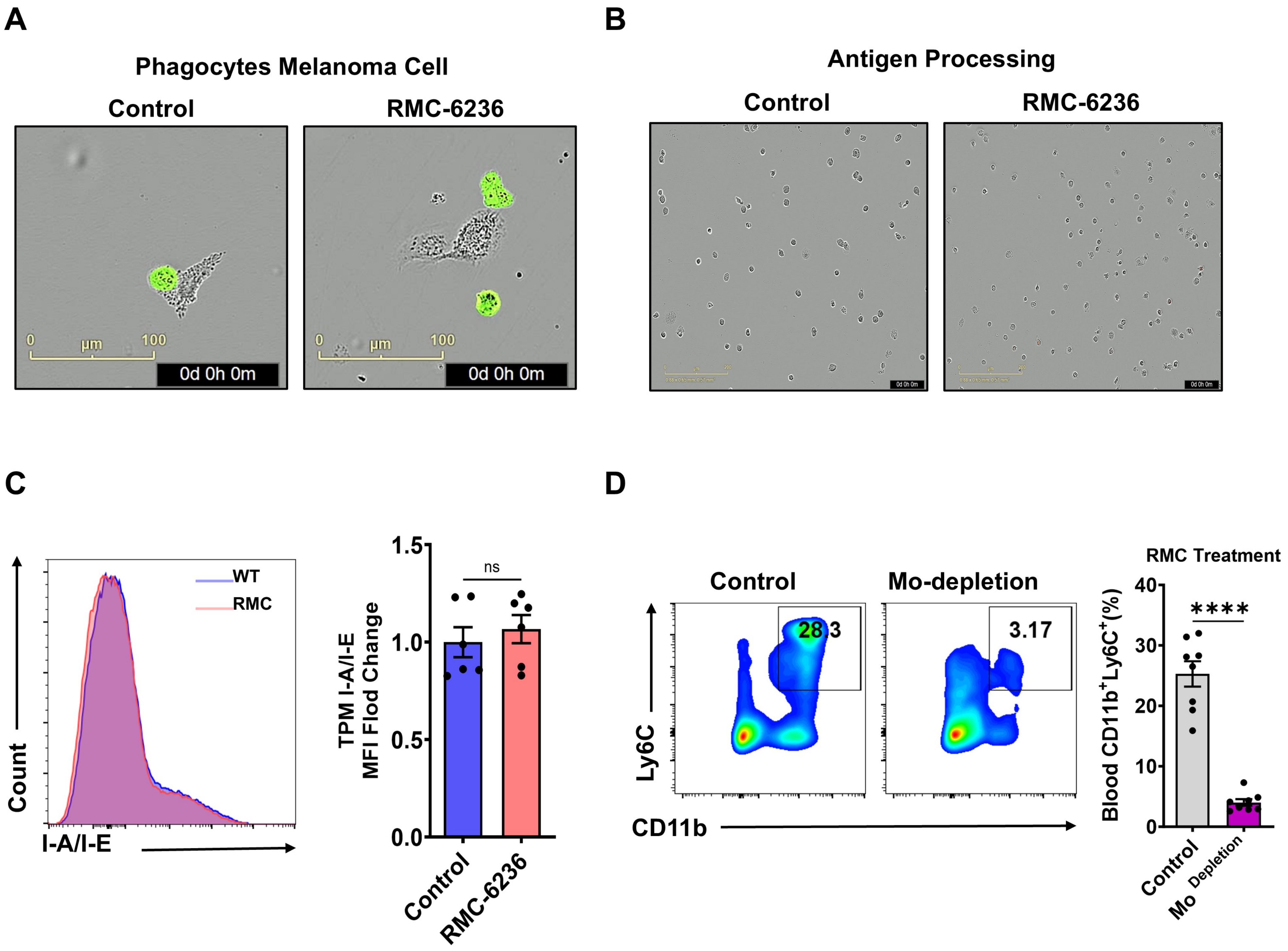

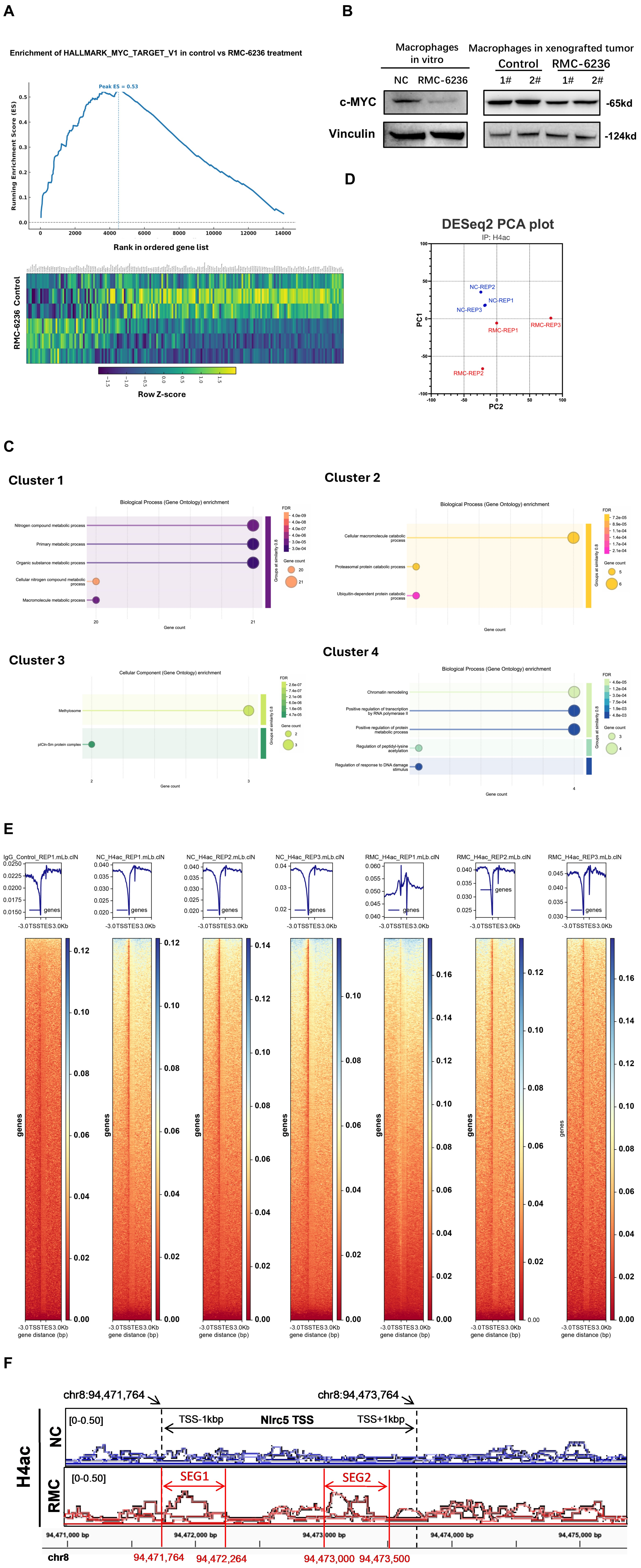

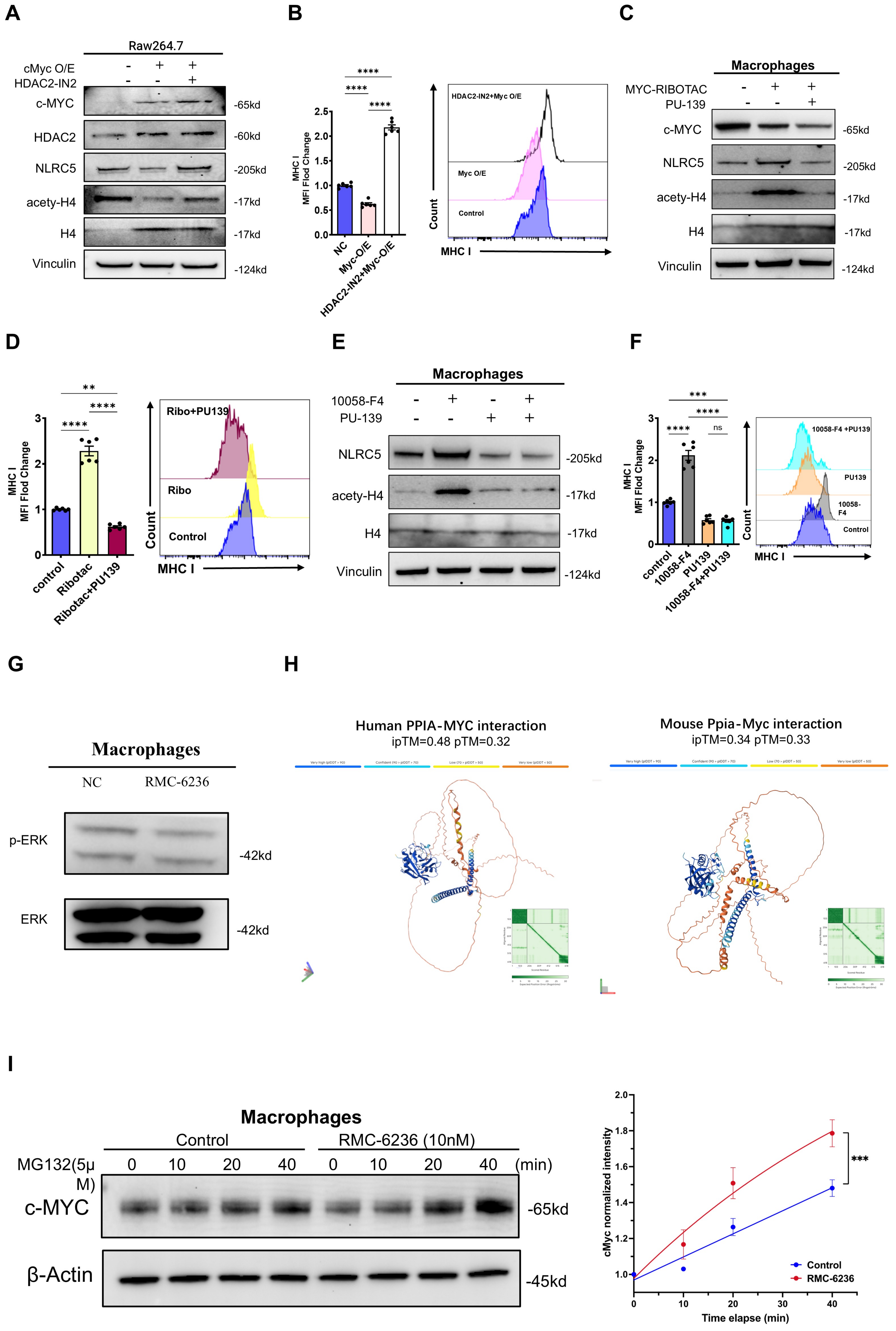
